# Parent-Child Autonomic Synchrony During Vicarious Extinction Learning in Pediatric PTSD

**DOI:** 10.1101/2022.01.12.476063

**Authors:** Grace C. George, Sara A. Heyn, Shuka Konishi, Marie-France Marin, Ryan J. Herringa

## Abstract

Children learn about threat and safety in their environment in part from their caregivers, a process which may be disrupted in child psychopathology. This transmission may be seen through biological measures like peripheral nervous system outputs such as skin conductance (SCR). Fear learning deficits have been observed in fear-related disorders like PTSD but have received little study in terms of parent-child learning transmission. We used a vicarious fear extinction paradigm to examine whether biological synchrony (SCR and heart rate variability [HRV]) is a potential mechanism by which children learn safety cues from their parents. In this pilot study, 16 dyads (PTSD n=11, typically developing [TD] n=5) underwent a vicarious fear extinction paradigm. We used cross-recurrence quantification analysis (CRQA) to assess SCR and HRV synchrony between parent-child dyads. We then used a linear model to examine group differences between PTSD dyads and TD dyads. For SCR, a significant group difference (p=.037) indicated that TD dyads had higher SCR synchrony compared to PTSD dyads. For HRV, there were no group differences between PTSD and TD dyads (p=.325). In exploratory analyses, increased synchrony was related to an overall decreased autonomic reactivity during recall of fear (p=0.032). These results suggest that SCR synchrony, but not HRV, may be a potential mechanism that allows for fear and safety learning in youth. While these data are preliminary, they provide novel insights on how disruptions in vicarious extinction learning may contribute to fear-related disorders in youth. Furthermore, this study suggests novel approaches to studying dyadic trauma-focused therapies which critically rely on parental coaching to model appropriate fear responses to help their child to recover from trauma.

**Significance Statement:** This study provides evidence that biological synchrony is a potential mechanism through which youth learn threat and safety cues through their parents. We found that youth with PTSD have lower synchrony with their caregiver, and that synchrony was related to decreased overall autonomic reactivity, suggesting that synchrony may be indicative of overall safety learning. Further, decreased synchrony during fear extinction may potentially underlie the etiology of fear related disorders such as PTSD. These novel approaches may improve our understanding of dyadic trauma-focused therapies which critically rely on parental coaching to model appropriate fear responses to help their child to recover from trauma.

## Introduction

Children’s ability to learn emotional content from their caregivers is an important aspect of child development (Debiec & Olsson, 2017). However, the transmission of emotional content may be altered in settings of parent or child psychopathology, as well as trauma, contributing to the emergence and/or persistence of fear-related disorders in youth (e.g. anxiety, PTSD) (Silvers et al., 2020; Stenson et al., 2020). In a clinical setting, especially for Trauma-Focused Cognitive Behavioral Therapy (TF-CBT) which utilizes dyadic treatment to help youth with trauma disorders, heightened transmission of threat signals from caregiver to child may hinder youth from reducing their symptom load (Golkar et al., 2016). Thus, it is critical to understand how children vicariously learn from their caregivers to better help treatment of fear-related disorders.

Fear extinction learning has been widely used to understand fear-related disorders like PTSD in adults, but with less study in children (Herringa et al., 2013; Milad et al., 2014). Previous studies in humans, with support from rodent models, have indicated alterations in fear learning like enhanced acquisition and impaired extinction in those with anxiety disorders (for a review see Milad et al., 2014). Of particular salience for youth, altered fear learning may partly be due to youths difficulty to learn fear and safety by observation, or vicarious learning, as this learning has been shown to influence a child’s normal fear development (Rachman, 1977). During fear acquisition, one study found that anxious children had increased reactivity to fear acquisition, demonstrating that psychopathology may influence vicarious fear learning (Bilodeau-Houle et al., 2020). While fear learning may be enhanced in youth with trauma, fear extinction or extinction recall may also be disrupted (Marusak et al., 2020). Therefore, it is important to study fear extinction and recall, in addition to acquisition, to fully understand the impact of vicarious fear learning on psychopathology in youth.

One mechanism through which vicarious extinction may occur is through parent-child physiological synchrony. Synchrony is the temporally-matched coordination of responses between two people (Feldman, 2012). For parent-child dyads, synchrony is a critical method of learnt emotion regulation in children and a way to foster healthy attachments (Davis et al., 2017). Physiological synchrony uses peripheral nervous system methods like skin conductance response (SCR) or heart rate variability (HRV) to evaluate the degree to which dyads are coupled (Feldman, 2012). Youth or parent trauma leads to differences in physiological, or autonomic, synchrony but how these variations affect real world behaviors like fear learning is still unknown (Feldman, 2007; Motsan et al., 2020). Understanding the biological mechanism behind vicarious learning is crucial for understanding transmission of fear and safety cues between dyads, especially in those with fear-related disorders like PTSD.

To address these knowledge gaps, this pilot study examined physiological synchrony during vicarious extinction learning in youth with PTSD compared to typically developing (TD) non-trauma exposed youth. During this paradigm, youth went through both direct and vicarious extinction, which included watching their parent undergo direct extinction. Preliminary evidence from the paradigm validation study suggests that youth with PTSD have increased SCR during vicarious fear learning compared to TD youth (Heyn et al., 2022). This demonstrates that there maybe be a biological mechanism at play when youth are learning fear and safety cues from their caregivers, and potentially this is disrupted in youth with PTSD. Here, we expand on this finding to understand if biological synchrony during vicarious fear extinction is altered in youth with PTSD.

We will be using two biological metrics to assess fear learning: SCR and HRV. SCR is a widely used measure of physiological arousal and is one of the most common biological metrics of fear condition and extinction (Faghih et al., 2015). Regulation of fear responses is linked to HRV as a reflection of amygdala (Schiller et al., 2008). We assessed SCR and HRV group differences in biological synchrony during vicarious learning. Then, for any significant group differences, we analyzed if synchrony was related to psychophysiology during recall and dyadic psychopathology symptoms.

We predicted that PTSD dyads will have lower biological synchrony compared to their typically developing counterparts. Additionally, we predicted that PTSD related symptoms will be associated to synchrony. Lastly, in exploratory analyses, we expected that synchrony will predict psychophysiology during recall, but only for the vicarious conditioned stimulus.

## Materials and Methods

In this pilot study, we recruited 16 parent-child dyads with youth ranging from ages 7-17 years. 11 of those dyads included a child with PTSD (10 female) and five (4 female) that were typically developing (TD). Exclusion criteria for our youth participants included past or present brain injury, unstable or sever medical conditions, substance abuse, acute suicidality, or ongoing abuse. Each parent-child dyad was assessed for past and current psychopathology diagnosis, including PTSD status, using the Mini-International Neuropsychiatric Interview Screen (MINI; Sheehan et al., 1998). Further psychopathology questionnaires for the child included the Mood and Feelings Questionnaire (MFQ) for depression, the Screen for Child Anxiety-Related Emotional Disorder (SCARED) for anxiety, and the UCLA PTSD Reaction Index (PTSD-RI) for PTSD symptoms (Birmaher et al., 1997; Costello & Angold, 1988; Steinberg et al., 2004). The PTSD-RI for the DSM-V and DSM-IV were given to different participants, and only the congruent items between the two were used for analysis (Cheng et al., 2021).

### Experimental Design

In the current study, parent-child dyads underwent a three-day vicarious and direct fear learning paradigm. We used an adaptation from Milad and colleagues (Milad et al., 2007) and is described in detail in Heyn et al., 2022. Briefly, each dyad completed a fear learning paradigm separately. On day one, both parent and child were conditioned to two colored stimuli (CS+), while the remaining stimulus was left unpaired (CS−). On the second day, both parent and child went through extinction training. During extinction training for the parent, one CS+ was extinguished while the other CS+ was left unextinguished. For the child, both CS+s were extinguished, one by direct extinction learning (CS+D) and the other vicariously extinguished by watching their parent (CS+V). Day three consisted of recall for the dyads. All three task days were approximately 24 hours apart. For the unconditioned stimulus, we used tactile electrodermal stimulation. Each participant was allowed to manually select their level of stimulation. No participants dropped out of the pilot study due to intolerance of the stimulation. Further discussion of experimental design can be found in Heyn et al. 2022.

During vicarious learning, we measured SCR, heart rate (HR), and respiration of each dyad. For the synchrony analyses, we used SCR and HRV. For both SCR and HRV, we cut each timeseries at the beginning of the first fixation to the beginning of the last fixation. SCR analyses include a low-pass filter of 1 Hz and down sampling to 8 Hz using Ledalab (Benedek & Kaernbach, 2010). HRV analyses used HR and respiration to create time and frequency domains using MindWare software (MindWare Technologies Inc., Westerville, OH).

### Statistical Analysis

Statistical analyses were performed in RStudio (RSTudio Team, 2012). For each synchrony analysis, we used cross-recurrence quantification analysis (CRQA) using the R package *crqa* (Coco & Dale, 2014). In brief, CRQA captures recurring properties and patterns of two distinct time series. Increased CRQA measures, or synchrony, indicates that the two time-series, for example parent-child SCR, resemble each other or mimic each other over time. We followed Pärnamets et al. (2019) parameters for the CRQA analysis. We then picked three metrics (Determinism, Entropy, and Laminarity) that were highly correlated r > .90 and conducted a Principal Component Analysis with varimax rotation to find a single composite score of synchrony using the *psych* package in R. For our main analyses, we conducted a linear regression for group (TD vs PTSD) while covarying for child age and sex. We then conducted post-hoc analyses on significant group differences. To test specificity to vicarious learning, we also examined group differences on synchrony when both the child and parent were undergoing direct extinction. We then conducted Pearson correlations between child symptoms from the PTSD-RI, MFQ, SCARED, and synchrony with FDR correction. We further covaried for parent age and lifetime or current psychopathology diagnosis of the parent on the group differences. Lastly, we conducted three exploratory repeated measures analyses to understand if synchrony was related to fear extinction outcomes measures. First, to assess if synchrony was related to fear learning responses, we conducted a repeated measures regression to see if synchrony and CS type predicted extinction retention index (ERI). ERI was calculated by taking the average SCR of the first 2 recall trials and dividing it by the highest SCR during the conditioning trials. Second, examined whether synchrony was related to child SCR responses during early recall (first four trials) and if these responses were moderated by CS type. Lastly, we wanted to understand if synchrony was related to expectancy of the shocks during the first trial of recall and if this was moderated by CS type. All models, besides the correlations, were covaried for child age and sex and Z-scored. Due to the skew of the recall data, all recall SCR data was log transformed and then Z-scored.

## Results

Participant demographics can be found in Table 1. There was a significant SCR group synchrony difference between the PTSD and TD groups during vicarious extinction learning b =1.25, t(13) = 2.34, p = .037 and an effect size of η2 = .31 (Figure 1). There was no significant difference between groups during direct extinction for SCR synchrony b = −0.18, t(13) = .35, p=.62. For HRV, there was no group synchrony difference during vicarious extinction b=.40, t(13)= 1.03, p=.325, η2 = .08.

**Table 1:** Participant Demographics.

| Table 1: Participant Demographics |  |  |  |  |
| --- | --- | --- | --- | --- |
|  |  | Typically Developin<br>g | PTSD | Group Comparisons |
| Basic Demographic Variables |  |  |  |  |
| N Dyads |  | 5 | 11 |  |
| Child Sex (Female) | | 4 | 10 | $c^2(1, N = 16) = 0.37, p = 0.54$ |
| Parent Sex (Female) | | 5 | 11 | $c^2(1, N = 16) = 0.48, p = 0.49$ |
| Child Age | | 11.19 | 13.86 | $t(15) = -1.31, p = .21$ |
| Parent Age | | 44.74 | 41.71 | $t(14) = .75, p = .47$ |
| Did Parent Receive a Shock (No) | | 7 | 1 | $c^2(1, N = 15) = 1.64, p = 0.20$ |
| Race | White | 4 | 7 |  |
|  | African American | 0 | 1 |  |
|  | Asian | 0 | 0 |  |
|  | Two or More | 1 | 1 |  |
|  | Hispanic or Latino | 0 | 2 |  |
|  | Not Hispanic Latino | 5 | 7 |  |
|  | Not Provided | 0 | 2 |  |
| Parent Education Level | Some High School | 0 | 0 |  |
|  | High School Degree | 0 | 3 |  |
|  | Some College | 1 | 2 |  |
|  | College Degree | 1 | 3 |  |
|  | Graduate Degree | 3 | 0 |  |
|  | Not Provided | 0 | 3 |  |
| Trauma Variables |  |  |  |  |
| PTSD-RI Total | | 8.75(8.32) | 31.09 (9.47) | $t(13) = -3.88, p=.002$ |
| PTSD-RI Reexperiencing | | 2(2.92) | 9.09 (4.29) | $t(13) = -2.85, p=.014$ |
| PTSD-RI Avoidance | | 2.5(1.66) | 10.82(4.93) | $t(13) = -3.08, p = .009$ |
| PTSD-RI Hyperarousal | | 4.25(3.96) | 11.18(4.32) | $t(13) = -2.61, p = .022$ |
| MFQ |  | - | 19.1(8.97) |  |
| SCARED |  | - | 32.5(11.91) |  |
| Parent Current Diagnosis (Presence) | | 1 | 5 | $\chi^2(1, N = 13) = 2.24, p = 0.13$ |
| Parent Lifetime Diagnosis (Presence) | | 2 | 5 | $\chi^2(1, N = 13) = 0.63, p = 0.43$ |
| **Table 1: Full Sample Participant Demographics.** Both parent and youth groups did not differ significantly in sex or age. There was a significant difference between groups for the PTSD-RI but did not differ on parent diagnoses. |  |  |  |  |
| *Abbreviations:* PTSD, Posttraumatic Stress Disorder; PTSD-RI, PTSD-Reaction Index; MFQ, Mood and Feelings Questionnaire; SCARED, Screen for Child Anxiety-Related Mood Disorders. |  |  |  |  |

**Fig 1.**
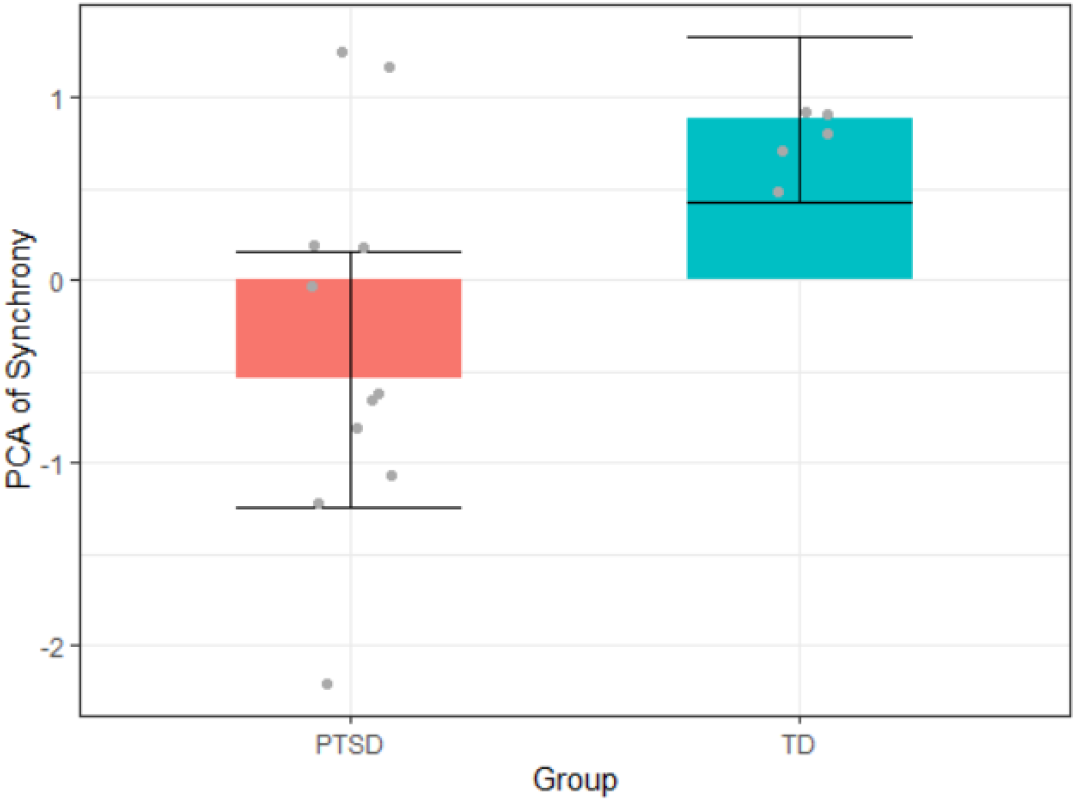
Group synchrony differences between PTSD and TD dyads. Using a linear model, we found a significant group difference between PTSD and TD dyads. Overall, TD dyads showed higher synchrony compared to PTSD dyads.

Due to the group differences, we also tested whether synchrony was related to child symptoms of depression, anxiety, or PTSD. There were no correlations between synchrony and any of the child symptom measures p > .27 (Table 1). Additionally, group differences remained marginally (significant) after covarying for parent age (b= 0.72, t(12) =1.27, p=.23), parent lifetime diagnosis (b= 1.06, t(12) = 1.01, p=.08), and parent current diagnosis (b=1.04, t(12) = 1.9, p=.09), suggesting group differences in synchrony were not attributable to these factors.

For the exploratory analyses, synchrony by CS type and the main effects did not significantly predict ERI, p>.19. For the second analysis, there was a significant main effect of vicarious SCR synchrony on recall SCR F(11,12) =4.62, p=0.032 indicating that synchrony during vicarious extinction learning was inversely related to SCR during recall (Figure 2). However, there was no significant interaction (p>.9) between synchrony and CS type on SCR during recall. For the third analysis, there was a significant CS type by synchrony interaction F(11,12) = 3.62, p=.02 for expectancy (Figure 3). Subsequently, we assessed if any of the CS type-synchrony slopes were significantly different from zero. The CS+D (t(11) = −.77, p=.46) and CS+V (t(11)= 1.28, p=.23) synchrony slopes were not significantly different from zero, but CS− was marginally different from zero (t(11) = −2.02, p=.068). Here, synchrony during vicarious extinction learning was inversely related to child stimulation expectancy for the CS− during extinction recall.

**Fig 2.**
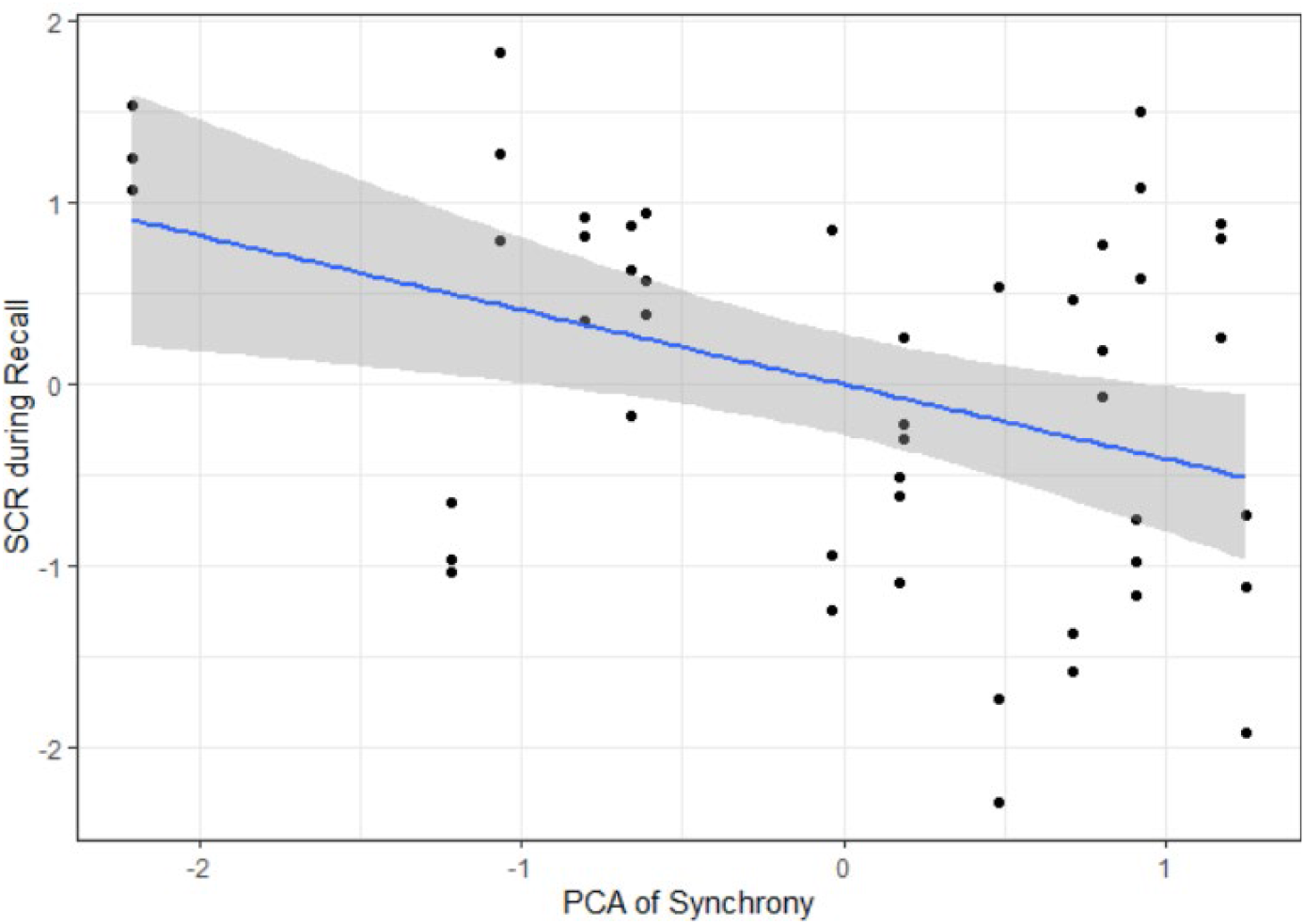
Synchrony is inversely related to child SCR during extinction recall. We found a significant main effect of synchrony on SCR during recall indicating that synchrony is related to overall decreases in arousal, but not CS specific decreases.

**Fig 3.**
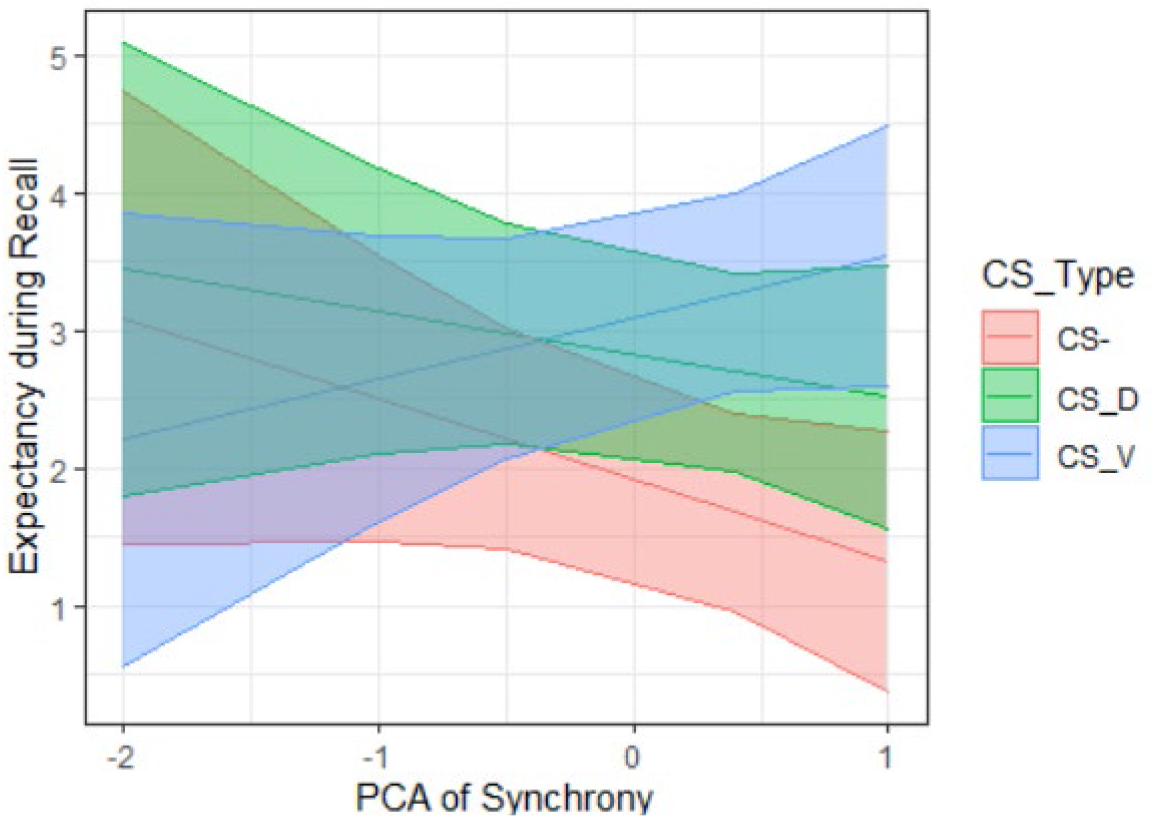
Significant CS type and synchrony interaction predicting expectancy. We found a significant interaction of CS type and synchrony on expectancy during recall. Only the CS− expectancy-synchrony slope was marginally significantly different from zero suggesting that synchrony may be a predictor of explicit safety instead of fear learning.

## Discussion

In this pilot study, we explored potential biological mechanisms that are related to vicarious fear and safety learning, specifically in a difficult group to recruit, youth with PTSD. We hypothesized that synchrony, or the coupling of two biological systems, in this case parent and youth psychophysiological outputs, is an important marker and mechanism of transmission of cues between each dyad and this may contribute to the expression of PTSD in youth. Overall, we found group differences in SCR synchrony between dyads that have youth with PTSD compared to typically developing dyads. In exploratory analyses, preliminary evidence showed that synchrony may lead, or be related to, SCR reactivity during recall indicating the possibility that dyads with higher synchrony during extinction learning have lower overall arousal during recall. Further, synchrony was moderated by CS type to predict expectancy. While both the CS+V and CS+D slopes were not different from zero, for the CS− (p=.068), increased synchrony was marginally related to less expectancy of a stimulation. This may indicate that increased synchrony is related to better learning of safety cues.

Overall, we found that youth with PTSD had lower SCR synchrony compared to their TD counterparts (Figure 1). However, we needed to account for the possibility that synchrony was not simply due to heritable patterns of extinction learning, but that it was specific to vicariously learning from (watching) the parent. To account for this, we conducted the same analysis with the child’s direct extinction SCRs and did not find this group difference, suggesting specificity of synchrony during extinction learning to vicarious learning.

While we predicted SCR and HRV to be significant in our main analyses, only SCR synchrony was significantly different between groups. Previous studies demonstrated that SCR synchrony was associated with greater threat learning and that increased SCR synchrony during fear acquisition was related to fear conditioned responses in parent-child dyads (Marin et al., 2020; Pärnamets et al., 2020). Adult PTSD studies found increased SCRs during recall suggesting a lack of fear extinction. However, studies of youth with trauma observed mixed results indicating that much is still unknown about how youth with psychopathology learn and extinguish fear (Garfinkel et al., 2014; Marusak et al., 2020; McLaughlin et al., 2016; Milad et al., 2009). For example, McLaughlin and colleagues found blunted SCRs to the CS+ and poor differentiation of CS types during conditioning and extinction in maltreated youth, while Marusak and colleagues found no differences in SCRs between TD and maltreated youth but did see behavioral differences of fear learning. The original pilot study specifically observed group differences in SCR during *vicarious* but not direct extinction learning (Heyn et al. 2022). Thus, social fear learning may represent an influential process in trauma-related disorders for youth, though further study is warranted. Our synchrony analyses strengthen that argument, as synchrony, which a known mechanism of learning (Davis et al., 2017), was blunted in dyads that had a child with PTSD.

Currently, there is less evidence to implicate HRV with synchrony and extinction learning. Previous studies examining HRV synchrony suggest relationships with positive attributes such as in higher levels of closeness, trust, and prosocial behaviors (Danyluck & Page-Gould, 2019; Goldstein et al., 1989). Individuals with social anxiety disorder (SAD) had difficulty in producing HRV synchrony in more intimate social contexts compared to individuals without SAD, thus leading to the decreased ability in developing relationships (Asher et al., 2021). However, these studies used general play or free-roam behavioral tasks instead of a structured fear extinction task. Further exploration of youth with PTSD and HRV is needed to understand how or if HRV is related to vicarious fear extinction.

For our exploratory analyses, we examined whether synchrony could be used to predict specific outcomes of fear extinction learning - ERI, SCR during recall, and expectancy of stimulation - and if this was moderated by CS type (CS+D, CS+V, and CS−). ERI is commonly used as a measure of extinction learning, as it takes into account baseline levels of autonomic reactivity during fear acquisition and compares it to outputs during recall (Milad et al., 2008). We did not see any significant main effects or interactions in this model. This may be due to the modest sample size, or more SCR variability in youth (Miller & Shields, 1980).

We conducted the second exploratory analysis to investigate the relationship between synchrony and CS type to SCR during recall. This analysis had a significant main effect of synchrony which showed increased synchrony being related to decreased SCR during recall, but not a significant synchrony by CS type interaction (Figure 2). It is possible that youth overall learned that this was safe environment from their parent leading to overall decreases in arousal. Synchrony could also be due to relationship quality between dyads. Relationship quality appears to be related to synchrony and therefore it is possible this is also related to overall lower arousal (Woody et al., 2016). In this pilot study, we did not collect parent-child relational measures, but future studies should consider adding these measures to parse why and how synchrony is related to overall arousal.

For our last exploratory analysis, we found a significant interaction between CS type and synchrony to predict expectancy of a stimulation, or the unconditioned stimulus (Figure 3). While the three CS types were significantly different from each other, we wanted to explore if any of the synchrony-CS relationships were different from zero. After running three linear models, only the CS− was marginally different from zero in slope. This, combined with our second exploratory analysis, indicates that synchrony may be preferentially related to safety and overall arousal as opposed to specific extinction of threat memory.

There are three main limitations that must be considered in this study. First, because of its nature as a pilot study, the sample size is modest and should be expanded to a larger population in future work. Therefore, these analyses should be interpreted with caution but may be useful to help guide future studies. Second, this study incorporated limited parent psychopathology measures. It is well known that parent psychopathology and other parental factors including parenting style and relationship affect the child’s propensity for psychopathology (Zhang et al., 2020). In this study we assessed parents with the MINI, which has individual diagnoses, but we used a binary of current or lifetime DSM-IV psychopathology diagnoses due to the modest sample size. In the future, it would be beneficial to utilize psychopathology symptom severity scores as well as presence of specific diagnoses in a larger sample. Further, it would be useful to see if parent styles or parent-child relationships are related to synchrony during extinction learning. Lastly, this study did not include trauma-exposed controls. Inclusion of this group in future work could help account for general effects of trauma and its relationship to synchrony and vicarious extinction learning. This also may help to answer potential questions about whether parenting styles or relationships may buffer the effect of trauma through synchrony.

In summary, this preliminary study provides novel insights into potential biological mechanisms underlying vicarious extinction learning in youth with PTSD and how this may contribute to trauma disorders in youth. Decreased synchrony in youth with PTSD could represent a biomarker and could contribute to maintenance of fear-related disorders and continued symptoms. Additionally, increased synchrony related to overall decreases in arousal indicating less reactivity during fear extinction recall overall. Learning proper safety and fear cues from safe caregivers is crucial to navigating the world effectively. Expansion of this work in future studies may help to unravel how inappropriate modeling of fear or altered vicarious learning in youth may contribute to the emergence and/or persistence of fear-related disorders in youth. If substantiated, such findings also carry notable implications for dyadic therapies, highlighting potential biomarkers that could be used to identify points of altered threat and safety transmission in the family system.

## ACKNOWLEDGEMENTS

We would like to thank the Brave Research Lab with their help in recruitment and data collection for this study and their feedback on the results. We also want to extend our genuine gratitude to the families and youth who have participated in our study.

